# B cell defects observed in *Nod2* knockout mice are a consequence of a *Dock2* mutation frequently found in inbred strains

**DOI:** 10.1101/242586

**Authors:** Serre-Yu Wong, Maryaline Coffre, Deepshika Ramanan, Luis E. Gomez, Lauren Peters, Eric Schadt, Sergei Koralov, Ken Cadwell

**Affiliations:** Kimmel Center for Biology and Medicine at the Skirball Institute, New York University School of Medicine, New York, NY 10016, USA.; Henry D. Janowitz Division of Gastroenterology, Icahn School of Medicine at Mount Sinai, New York, NY 10029, USA.; Department of Pathology, New York University School of Medicine, New York, NY 10016; Department of Genetics and Genomic Sciences, Icahn School of Medicine at Mount Sinai, New York, New York, 10029, USA.; Icahn Institute for Genomics and Multiscale Biology, Icahn School of Medicine at Mount Sinai, New York, New York, 10029, USA.; Sema4, a Mount Sinai venture, Stamford, Connecticut, 06902, USA.; Department of Microbiology, New York University School of Medicine, New York, NY 10016, USA.

## Abstract

Phenotypic differences among substrains of laboratory mice due to spontaneous mutations or pre-existing genetic variation confound the interpretation of targeted mutagenesis experiments, and contribute to challenges with reproducibility across institutions. Notably, C57BL/6NHsd mice and gene-targeted mice that have been backcrossed to this substrain have been reported to harbor a duplication in exons 28 and 29 of *Dock2*. Here, we demonstrate the presence of this *Dock2* variant in the widely used *Nod2^−/−^* mice. NOD2 is a cytosolic innate immune receptor that has been the subject of intense investigation due to its association with inflammatory bowel disease (IBD) susceptibility. Consistent with a role of NOD2 in an immunological disorder, *Nod2*^−/−^ mice bred at our institution displayed multiple B cell defects including deficiencies in recirculating B cells, marginal zone B cells and B1a cells. However, we found that these effects are due to the *Dock2* variant and are independent of *Nod2* deletion. Despite originating from the same gene-targeted founder mice, *Nod2^−/−^* mice from another source did not harbor the *Dock2* variant or B cell defects. Finally, we show that *Dock2^−/−^* mice display the same B cell defects as mice harboring the *Dock2* variant, confirming that the variant is a loss-of-function mutation and is sufficient to explain the alterations to the B cell compartment observed in *Nod2^−/−^* mice. Our findings highlight the effects of confounding mutations from widely-used inbred strains on gene-targeted mice and reveal new functions of DOCK2 in B cells.

## INTRODUCTION

Inbred laboratory mice are essential animal models that facilitate comparison of experimental outcomes observed across institutions. They are commonly used to identify the function of a gene product following a targeted mutagenesis event such as a gene knockout or the expression of a transgene. However, spontaneous mutations, that eventually contribute to genetic drift, can arise from maintaining breeding stocks separately at different vendors and institutions, and the effect of genetic variability among a given inbred strain is rarely considered. These concerns may have contributed to the recent controversy regarding reproducibility of findings using preclinical mouse models (1).

Among inbred mice, C57BL/6 is the most commonly used genetic background in immunology research. Although C57BL/6 substrains are known to harbor genetic differences (2), they are often treated as equal and are not distinguished in publications. The discovery of a mutation within the coding region of *dedicator of cytokinesis 2* (*Dock2*) in certain gene-targeted C57BL/6 lines highlights this issue. DOCK2 mediates actin polymerization and intracellular signaling as a guanine nucleotide exchange factor (GEF) for the small GTPase Rac (3). Consistent with its expression in the hematopoietic compartment, *DOCK2* mutations have been identified in immunocompromised children who display early onset invasive infections (4). Also, *Dock2^−/−^* mice display several immune defects including loss of marginal zone (MZ) B cells and impaired migration of lymphocytes, neutrophils, and plasmacytoid dendritic cells (pDCs) (3, 5–7). Similar immune defects observed in mice with other targeted mutations have now been attributed to a spontaneous mutation in *Dock2*. Loss of MZ B cells and reduced type I interferon (IFN) production by pDCs in *Irf5^−/−^* mice were shown to be independent of *Irf5* deletion and instead due to a duplication of exons 28 and 29 of *Dock2* that leads to reduced *Dock2* expression (8). Likewise, defects in B cell development observed in a subset of mice deficient in sialic acid acetyl esterase (*Siae^−/−^*) were due to a similar exon duplication in *Dock2*. This mutation in *Dock2* was traced to the C57BL/6NHsD substrain that was used to backcross the *Siae^−/−^* line onto the C57BL/6 background (9). Nonetheless, not all the B cell defects in these mutant mice were a consequence of the defective *Dock2* allele, as another study noted that differences in IgG isotype switching were found in *Irf5^−/−^* mice with and without the *Dock2* variant (10). Decreased *Dock2* expression has also been suggested to explain why a subset of mice deficient in the inflammasome adaptor ASC (*Pycard^−/−^*) display defects in antigen presentation and lymphocyte migration, although it is unclear whether these mice harbor the same duplication event as *Irf5^−/−^* mice (11).

Here, we demonstrate that B cell defects in *Nod2^−/−^* mice are due to *Dock2* mutation and independent of *Nod2* deficiency. NOD2 (nucleotide-binding oligomerization domain-containing protein 2) is a cytosolic pattern recognition receptor best known for controlling an antimicrobial gene expression program in response to peptidoglycan (12). Loss of function mutations in NOD2 are among the strongest susceptibility factors for Crohn’s disease, a major type of inflammatory bowel disease (IBD) characterized by chronic relapsing inflammation of the gastrointestinal tract (13). Inconsistent results obtained with *Nod2* mutant mice have created a major barrier to understanding the role of NOD2 in Crohn’s disease. *Nod2^−/−^* mice were originally shown to display defective defensin expression by Paneth cells (14), antimicrobial epithelial cells in the small intestine (15). However, this defect was not observed in commercially available *Nod2^−/−^* mice that were backcrossed onto the C57BL/6J background (16). Also, early findings demonstrating increased cytokine production and a T cell-intrinsic function in *Nod2* mutant mice were not reproduced in subsequent studies, potentially due to the presence of unintended mutations in the original mice that were characterized (17–22). Variation in the microbiota can also explain disparate results. *Helicobacter* species that are eradicated in some animal facilities induce an enhanced Th1 response in *Nod2^−/−^* mice that leads to inflammatory lesions in the small intestine (23). Although *Nod2^−/−^* mice are susceptible to colonization by *Bacteroides* species, control wild-type (WT) mice co-housed with *Nod2^−/−^* mice acquire a similar microbiota (24–26). We previously demonstrated that a particular *Bacteroides* species that is not present in commercially available *Nod2^−/−^* mice, *Bacteroides vulgatus*, mediates goblet cell defects in *Nod2^−/−^* mice and not WT mice raised in our vivarium (24, 27). Thus, genetic background and microbiota composition have profound influence on results obtained with *Nod2* mutant mice.

In this study, we identify deficiencies in populations of recirculating B cells in the bone marrow, MZ B cells, and splenic and peritoneal B1a B cells in *Nod2^−/−^* mice. These B cell defects were not present in mice deficient in the NOD2 signaling adaptor RIP2 (*Rip2*^−/−^) or a second *Nod2^−/−^* line that we acquired. We found that differences in phenotype were driven by the presence of the aforementioned *Dock2* mutation. Importantly, we demonstrate that independently generated *Dock2^−/−^* mice display similar B cell defects. All together, these findings reveal new functions of DOCK2 and show that certain lymphocyte-defects observed in *Nod2^−/−^* mice are independent of NOD2 function.

## MATERIALS AND METHODS

### Mice

*Nod2^−/−^* mice backcrossed to the C57BL/6 background for at least 12 generations were previously described (28). These mice were imported to Washington University School of Medicine and subsequently rederived into the animal facility at NYU School of Medicine where they have been maintained until present. *Nod2^−/−^* mice (*Nod2^−/−^Jax* mice) derived from the original gene targeting experiment (14) were obtained from The Jackson Laboratory and bred on-site. Wild-type C57BL/6J mice were purchased from The Jackson Laboratory and bred on-site. *Rip2^−/−^* mice bred on-site were previously described (24, 29). All on-site animals were maintained in a specific pathogen-free facility at the New York University School of Medicine. Experiments were approved by the Institutional Animal Care and Use Committee of the New York Unversity School of Medicine. *Dock2*^−/−^ mice and wild-type littermates used for experiments were generated from *Dock2*^+/−^ breeders and were a gift of Dr. Yoshinori Fukui of Kyoto University (3). These mice were bred and housed at a specific pathogen-free facility at Charles River Laboratories. Experiments were approved by the IACUC of the Icahn School of Medicine at Mount Sinai.

### Bone marrow chimeras

Bone marrow chimeras were generated by irradiating 8-week-old female *Rag1^−/−^* mice (1,100 CGy once) followed by intravenous injection of 5 × 10^6^ T-cell-depleted bone marrow cells mixed in a 1:1 ratio from donor WT and *Nod2^−/−^* female mice. Recipient *Rag1^−/−^* mice were treated with antibiotics in their drinking water one week prior to irradiation.

### Flow cytometry

Single cell suspensions from spleen, bone marrow, and cells from the peritoneal cavity were stained for cell surface markers. FACS analysis was performed on an LSR Fortessa cytometer. Antibodies to the following cell surface markers were used: anti-B220, CD19, CD25, c-kit, AA4.1 (CD93), CD21, CD23, CD3, CD5 (eBioscience), CD1d (Biolegend), and, IgM (μ chain specific) (Jackson ImmunoResearch). Data were analyzed using FlowJo software. Events are gated on total lymphocytes and singlets. Further gating is described in the figure legends when appropriate.

### PCR

Genomic DNA from mouse tails was harvested by NaOH extraction and analyzed for the presence of the *Dock2* duplication by PCR as previously described (9, 30). PCR primers used for the *Dock2* duplication were ln29.4F 5’-GACCTTATGAGGTGGAACCACAACC-3’ and lnR22.3.1R 5’-GATCCAAAGATTCCCTACAGCTCCAC-3’. PCR primers for the internal control (CD19) were oIMR1589 5’-CCTCTCCCTGTCTCCTTCCT-3’ and oIMR1590 5’-TGGTCTGAGACATTGACAATCA-3’.

### Statistical analysis

Statistical analysis was performed using GraphPadPrism 7. Differences in median values were analyzed by Mann-Whitney U test to compare two groups.

## RESULTS

### *Nod2^−/−^* mice are deficient in several B cell subsets

*Nod2^−/−^* mice are susceptible to disease caused by intestinal pathogens and pro-inflammatory members of the microbiota (23, 24, 26, 31). Given the fundamental role of B cells in mucosal immunity, and the role of antibodies in intestinal homeostasis and immune exclusion, we performed a broad flow cytometry-based analyses of B cell populations in the bone marrow and spleen of *Nod2^−/−^* mice to evaluate the contribution of these cells to colonization resistance. The *Nod2^−/−^* mice maintained in our animal facility were originally generated through gene targeting of 129/SvJ ES cells and backrossed over 10 generations onto the C57BL/6 background (see Methods). We found that these *Nod2^−/−^* mice lacked recirculating B cells defined as B220hi IgM+ cells in the bone marrow, despite displaying similar proportions of pro- and pre-B cells as well as immature B lymphocytes compared to wild-type (WT) C57BL/6J controls (Fig 1A, B). *Nod2^−/−^* mice also displayed marked splenocytopenia with a reduction in total B220+CD19+ B cell numbers compared to WT controls (Fig. 1C). Although the follicular B cell compartment remained intact, MZ B cells were entirely absent in the spleen of *Nod2^−/−^* animals (Fig. 1D).

**Figure 1.**
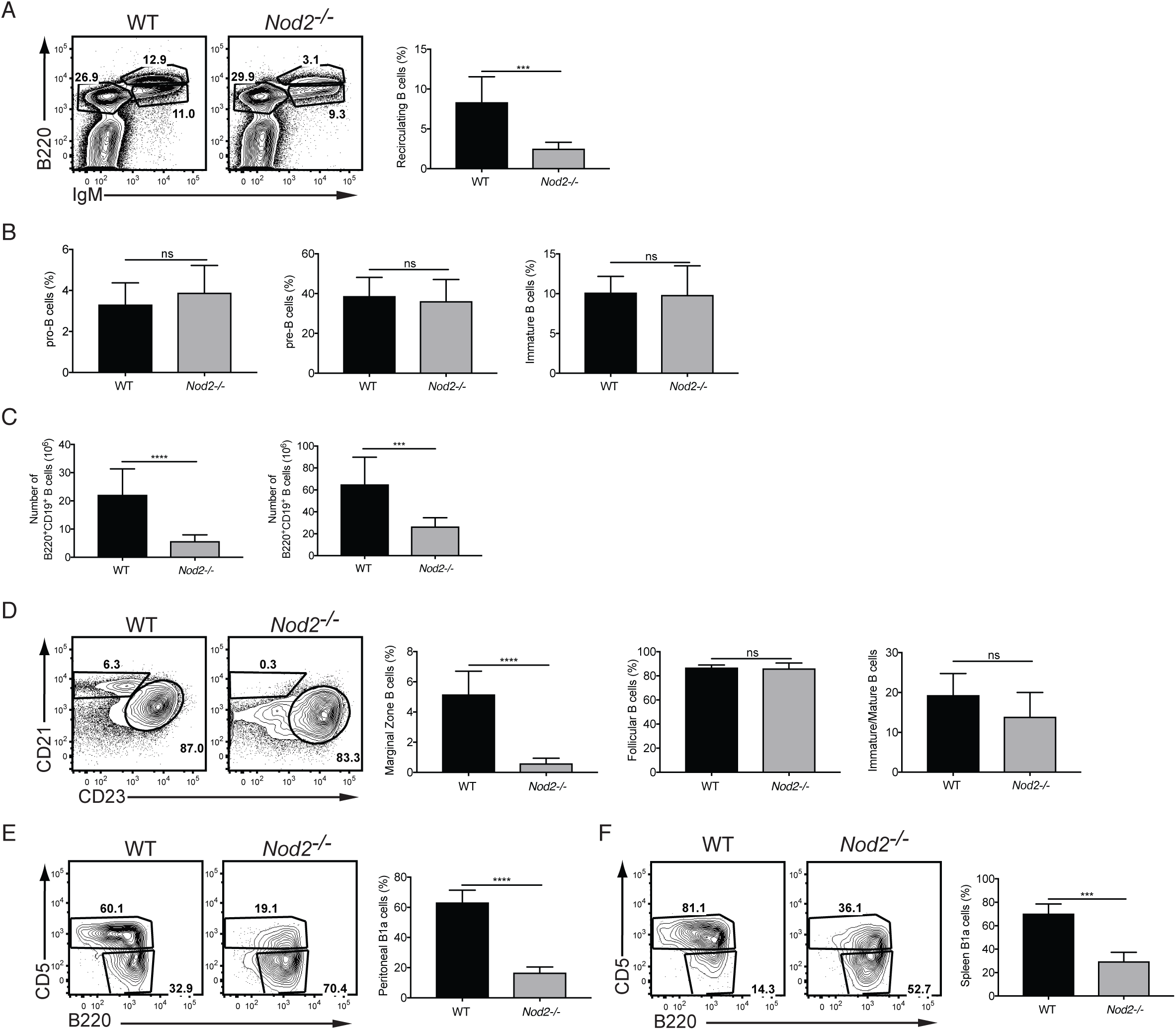
*Nod2^−/−^* mice have defects in recirculating B cells, marginal zone B cells, and B1a cells. Comparison of B cell populations in WT (*n* = 9) and *Nod2^−/−^* (*n* = 9) mice. **(A)** Representative plot and percentage of bone marrow recirculating B cells (B220high IgM+). **(B)** Percentages of pro-B, and pre-B, and immature B cells in the bone marrow. **(C)** Total cell numbers and proportion of B220+ CD19+ B cells in the spleen. **(D)** Representative FACS plot and percentages of marginal zone (CD21high CD23low) and follicular (CD21low CD23high) B cells in the spleen. Populations are gated on B220+ CD19+ AA4.1-mature B cells. **(E-F)** Representative plot and percentages of B1a cells (B220low, CD19+ CD5+) from peritoneal lavage (D) and in spleen (E). Data is representative of greater than five independent experiments. ns *p* > 0.05, \*\*\**p* ≤ 0.001, and \*\*\*\**p* ≤ 0.0001 by two-tailed Mann-Whitney.

B1 B cells participate in early immune responses as a first line of defense against a wide range of pathogens and are critical contributors to mucosal IgA responses (32, 33). Because of the shared signaling pathways downstream of Ig engagement between MZ and B1 B cells (34–36), defects in MZ development are often accompanied by a reduction in B1a B cells. We found that B1a B cells were reduced in the spleen and the peritoneal cavity of *Nod2^−/−^* compared to WT mice (Fig. 1E, F). These findings indicate that *Nod2^−/−^* mice maintained in our facility are deficient in several key B cell populations, specifically recirculating B cells in the bone marrow, MZ B cells in the spleen, and B1a B cells in the spleen and peritoneum.

### B cell defects in *Nod2^−/−^* mice are cell-autonomous

The lymphocyte-intrinsic function of NOD2 is controversial (17–22). To ask whether B cell defects are cell-autonomous or induced by extrinsic factors, we reconstituted *Rag1^−/−^* mice with mixed bone marrow from *Nod2^−/−^* mice (CD45.2+) and congenic CD45.1+ WT mice. We observed that the proportion of CD45.2+ cells in recipients were decreased when compared with CD45.1+ cells (Fig. 2A), suggesting a general defect in the development or maintenance of *Nod2^−/−^* leukocytes when forced to compete with WT cells. For subsequent analyses, the proportion of WT and *Nod2^−/−^* B cell populations were normalized to total B cells (B220+ cells) of the respective genotypes. The proportion of *Nod2^−/−^*-derived CD45.2+ recirculating B cells was reduced compared with WT-derived CD45.1+ cells (Fig. 2B). Analysis of spleens and peritoneal cells from the mixed bone marrow chimera mice revealed fewer *Nod2^−/−^*-derived MZ B cells and B1a cells (Fig. 2C-E). In contrast, although the total number of follicular B cells in the spleen were decreased due to the reduced reconstitution of the B cell compartment by *Nod2^−/−^*-derived cells, the relative proportion of *Nod2^−/−^* follicular B cells was similar to their WT-derived counterparts (Fig. 2C). Together, this data suggest a cell-autonomous mechanism for the loss of recirculating, MZ, and B1a B cells in the *Nod2^−/−^* mice.

**Figure 2.**
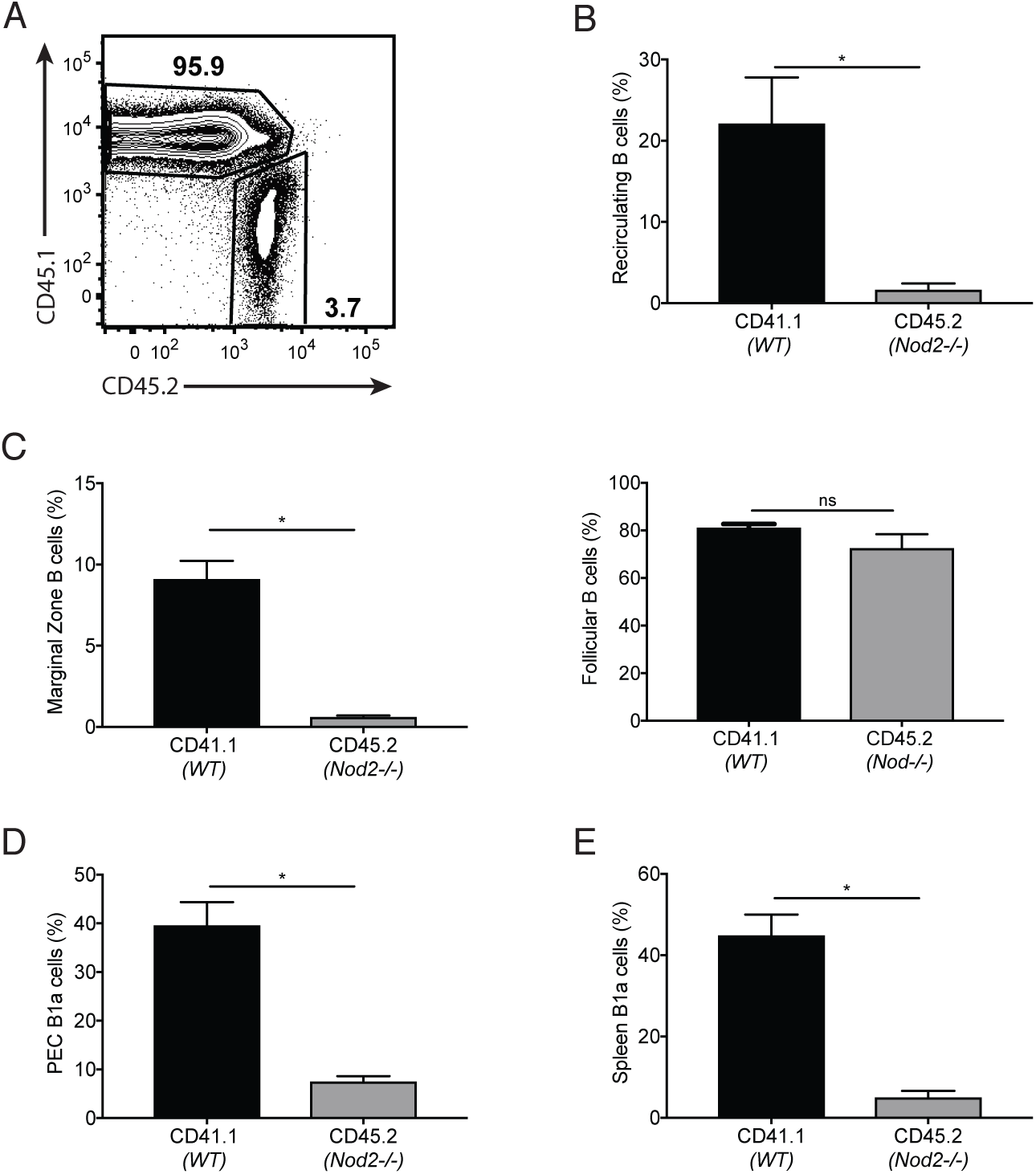
*Nod2^−/−^* cells are unable to efficiently reconstitute many of the B cell compartments. *Rag1*^−/−^ mice (*n* = 4) were reconstituted with mixed bone marrow from *Nod2^−/−^* (CD45.2+) and congenic CD45.1+ WT mice. **(A)** Representative plot of CD45.1+ and CD45.2+ mature B cells in the bone marrow. **(B)** Percentage of bone marrow recirculating B cells (B220high IgM+). Single cells were gated on CD45.1 or CD45.2 followed by IgM versus B220. The percentages of recirculation cells among CD45.1 or CD45.2 cells are shown. **(C)** Percentages of marginal zone (CD21high CD23low) and follicular (CD21low CD23high) B cells among CD45.1+ or CD45.2+ cells in the spleen. **(D-E)** Percentages of B1a cells (B220low, CD19+ CD5+) from peritoneal lavage (D) and spleen (E). CD19+ CD3-cells were gated to reveal CD45.1 and CD45.2 B cells followed by B1 gate (CD19+ B220 low) from which B1a cells were gated (CD5+ B220 low). Data is representative of two experiments. ns *p* > 0.05 and \**p* ≤ 0.05 by two-tailed Mann-Whitney.

### RIP2 deficiency does not recapitulate B cell defects observed in *Nod2^−/−^* mice

In the presence of the peptidoglycan derivative muramyl dipeptide (MDP), NOD2 engages receptor interacting protein 2 (RIP2) to initiate downstream NF-kB and MAPK signaling. In our previous study, we demonstrated that *Rip2^−/−^* mice phenocopied intestinal pathologies and susceptibility to alterations in the microbiota observed in *Nod2^−/−^* mice (24). We therefore asked whether disruption of RIP2 function would recapitulate the B cell defects seen in the *Nod2^−/−^* mice. Unexpectedly, recirculating B cells in the bone marrow and MZ B cells in the spleen of *Rip2^−/−^* mice were present in proportions similar to WT mice (Fig. 3A, B). Similarly, we did not observe a decrease in splenic or peritoneal B1a B cells in *Rip2^−/−^* mice (Fig. 3C, D). Thus, the B cell defects in the *Nod2^−/−^* mice are not observed in mice deficient in the signaling adaptor protein RIP2.

**Figure 3.**
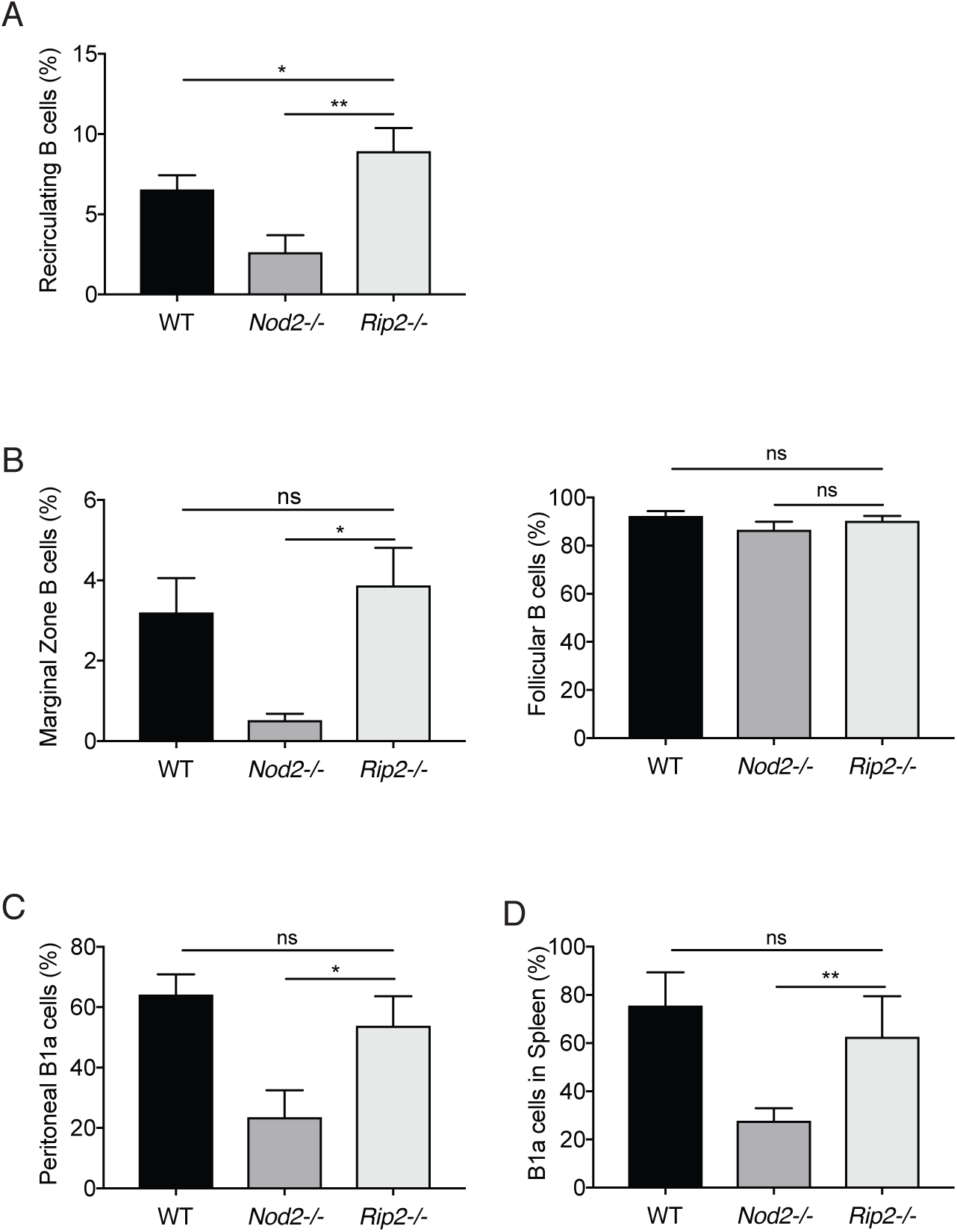
*Rip2*^−/−^ mice have normal recirculating B cell, marginal zone B cell, and B1a cell compartments. Comparison of B cell populations in WT (*n* = 4), *Nod2^−/−^* (*n* = 5), and *Rip2*^−/−^ (*n* = 6) mice. **(A)** Quantification of the total B cell number in the spleens of the animals. **(B)** Percentage of bone marrow recirculating B cells (B220high IgM+). **(C)** Percentages of marginal zone (CD21high CD23low) and follicular (CD21low CD23high) B cells in the spleen. Populations shown were gated on B220+ CD19+ AA4.1-mature B cells. **(D-E)** Percentages of B1a cells (B220 low, CD19+CD5+) from peritoneal lavage (C) and spleen (D). Data is representative of four independent experiments. ns *p* > 0.05, \**p ≤* 0.05, and \*\**p ≤* 0.01 by two-tailed Mann-Whitney.

### Commercially-available *Nod2^−/−^* mice do not show B cell defects

The observation that NOD2 may be functioning in a manner independent of RIP2 was unexpected. Thus, we sought to verify our findings by obtaining *Nod2^−/−^* mice from another source. *Nod2^−/−^* mice available for purchase from Jackson Laboratory were initially generated from the same gene targeted founders as the *Nod2^−/−^* mice maintained in our facility, but were backcrossed to the C57BL/6J background independently. These mice (*Nod2^−/−^Jax*) should be genetically identical to our *Nod2^−/−^* mice at the *Nod2* locus. However, whereas our *Nod2* mice have been backcrossed using an unspecified C57BL/6 substrain for 12 generations and have passed through multiple institutions (see Methods), the *Nod2^−/−^Jax* mice were deposited after six generations of backcrossing and backcrossed for one additional generation to C57BL/6J mice; the Jackson Laboratory reports that single nucleotide polymorphism (SNP) analysis shows that these mice are on a mixed C57BL/6J and C57BL/6N background. Analyses of bone marrow, spleen, and peritoneal cells showed that the *Nod2^−/−^Jax* mice did not harbor any of the B cell defects found in the *Nod2^−/−^* mice (Fig. 4). Thus, commercially available *Nod2^−/−^Jax* mice do not reproduce the reduction in B cell populations observed in our *Nod2^−/−^* colony.

**Figure 4.**
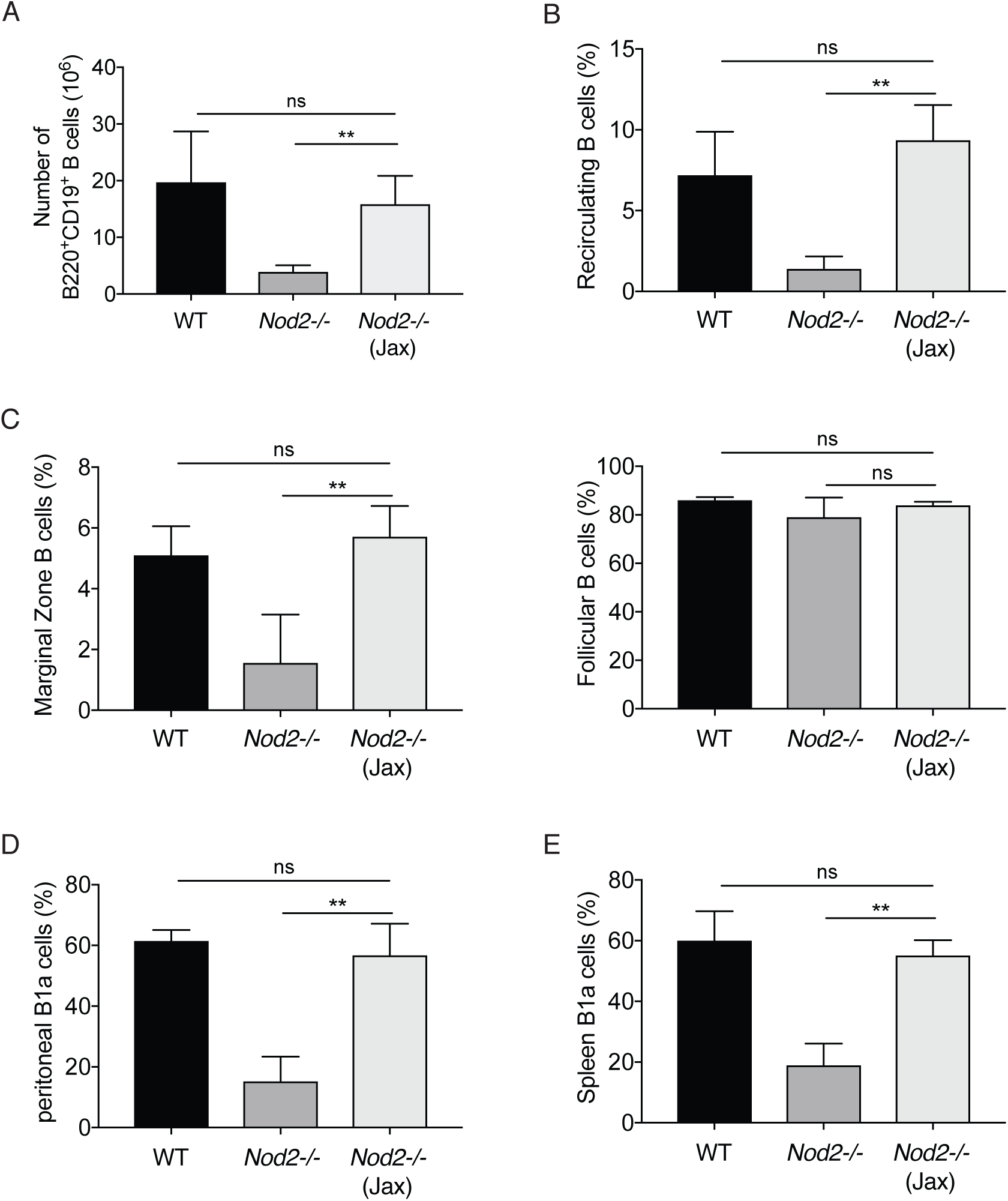
*Nod2*^−/−^ Jax mice have no defects in recirculating B cell, marginal zone B cell, and B1a cell compartments. Comparison of B cell populations in WT (*n* = 3), *Nod2^−/−^* (*n* = 5), and *Nod2^−/−^* Jax (*n* = 6) mice. (A) Percentage of bone marrow recirculating B cells (B220high IgM+). (B) Percentages of marginal zone (CD21high CD23low) and follicular (CD21low CD23high) B cells in the spleen. Population are gated on B220+ CD19+ AA4.1-mature B cells. (C-D) Percentages of B1a cells (B220 low, CD19+ CD5+) from peritoneal lavage (C) and spleens of mutant and WT mice (D). Data is representative of three independent experiments. ns *p* > 0.05 and \*\**p* ≤ 0.01 by two-tailed Mann-Whitney.

### B cell defects in *Nod2^−/−^* mice are associated with a mutation in *Dock2*

The disparity between the two *Nod2^−/−^* lines we observed could be due to an unknown genetic factor or a difference in an environmental variable, such as the microbiota. In our previous study, we demonstrated that antibiotics-treatment of *Nod2^−/−^* lines reversed intestinal pathologies (24). However, administration of antibiotics did not alter the number or proportion of B cell subsets in *Nod2^−/−^* mice maintained at our institution, and co-housing the two *Nod2^−/−^* lines did not transfer the phenotype in either direction (data not shown). Because the *Nod2^−/−^* animals in our colony were maintained as homozygous mice, we sought to eliminate the effect of the microbiota and avoid artifacts due to housing conditions by examining animals derived from an intercross between *Nod2^+/−^* mice. To generate littermate WT (*Nod2^+/+^*) and *Nod2^−/−^* mice for comparison, *Nod2^−/−^* mice from our colony were crossed to WT C57BL/6J mice and F1 *Nod2^+/−^* mice were bred to each other to generate an F2 generation (Fig. 5A). Suprisingly, one of eight *Nod2^+/−^* mice from the first F2 litter showed B cell defects similar to the F0 *Nod2^−/−^* mice (Fig. 5B). None of the other mice from that litter, including those with a *Nod2^−/−^* genotype, harbored the B cell defects. Although it is theoretically possible that a microbial factor that blocks the B cell defect was transferred from WT to *Nod2^−/−^* mice during these crosses, it is difficult to reconcile this possibility with our finding that one of the *Nod2^+/−^* mice displayed the B cell defect. Instead, these results suggested the presence of an additional genetic alteration in the *Nod2^−/−^* mice that is causing B cell abnormalities.

**Figure 5.**
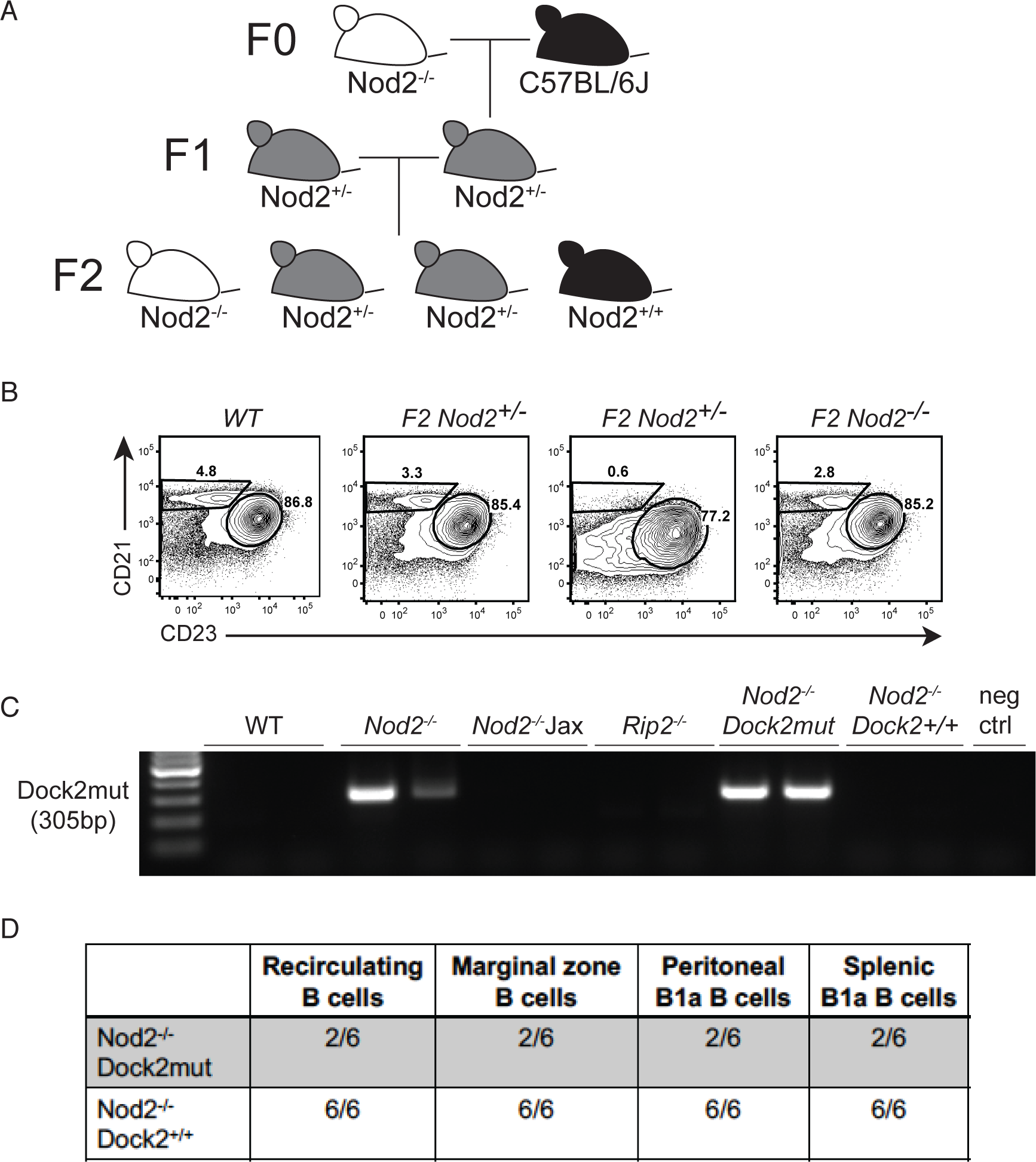
B cell defects in *Nod2^−/−^* mice are due to a mutation in *Dock2* and are independent of the *Nod2* deficiency. **(A)** Breeding scheme to segregate the *Nod2* targeted allele and to generate WT and *Nod2^−/−^* littermate controls. **(B)** Representative FACS plot showing percentages of marginal zone (CD21high CD23low) B cells in F2 generation littermates. **(C)** PCR for presence of *Dock2mut* in WT, *Nod2^−/−^, Nod2^−/−^* Jax, *Rip2^−/−^*, and F2 *Nod2^−/−^* mice. **(D)** Summary table showing number of mice with normal B cell populations as indicated among the total number of mice tested for the indicated genotypes. Data is representative of at least two independent experiments.

The splenocytopenia and MZ B cell deficiency we observed in our *Nod2^+/−^* mice are reminiscent of similar defects observed in *Siae^−/−^* mice, which were shown to be due to a duplication event involving exons 28 and 29 in *Dock2* (9). The *Dock2* mutation (*Dock2mut*) was shown to be present in the C57BL/6NHsD substrain and may have been introduced into gene-targeted mice during backcrossing (9). PCR analysis revealed the presence of the *Dock2mut* allele in our *Nod2^−/−^* mice, but not WT, *Nod2^−/−^Jax*, or *Rip2^−/−^* mice (Fig. 5C), suggesting that the presence of the *Dock2* exon duplication is responsible for the B cell defects we identified.

We then genotyped the F2 mice from the aforementioned F1 *Nod2^+^*^/−^ cross (Fig. 5A) for the presence of *Dock2mut*. The *Dock2* genotyping PCR, as previously described, only reveals the presence or absence of the duplication (9, 30) and does not indicate heterozygosity or homozygosity for the mutation. Thus, we relied on the analysis of the characteristic loss of MZ B cells (previously described to be a consequence of *Dock2* mutation (9)) to reveal which mice were homozygous for *Dock2mut*. We were able to separate the *Nod2* knockout allele from *Dock2mut* by analyzing F2 pups that were *Nod2^−/−^* with and without the presence of a *Dock2mut* allele. All six *Nod2^−/−^ Dock2^+/+^* mice tested had normal B cell compartments (Fig. 5D). In contrast, four *Nod2^−/−^* mice that presumably harbor two copies of the *Dock2mut* allele based on the loss of MZ B cells displayed reductions in recirculating B cells in the bone marrow and loss of peritoneal and splenic B1a cells (Fig. 5D). There were also two F2 *Nod2* mice that were positive for the *Dock2* mutation by PCR but did not show the B cell defects, likely because these animals carried one intact, WT copy of the *Dock2* allele.

### *Dock2*^−/−^ mice show B cell defects

The duplication in exons 28 and 29 lead to decreased expression of *Dock2* (8). Although the splenocytopenia and loss of MZ B cells have been described in other mouse lines that harbor the *Dock2mut* allele, we sought to determine whether loss of DOCK2 function is sufficient to cause the collection of B cell defects we observed. Therefore, we evaluated B cell subsets in *Dock2*^−/−^ mice generated by traditional gene-targeting (3). We found that *Dock2^−/−^* mice reproduced reductions in recirculating B cells in the bone marrow, MZ B cells in the spleen, and B1a B cells in the spleen and peritoneum observed in *Nod2^−/−^* mice harboring *Dock2^mut^* (Fig. 6). Therefore, DOCK2 function is essential for maintaining these key B cell populations.

**Figure 6.**
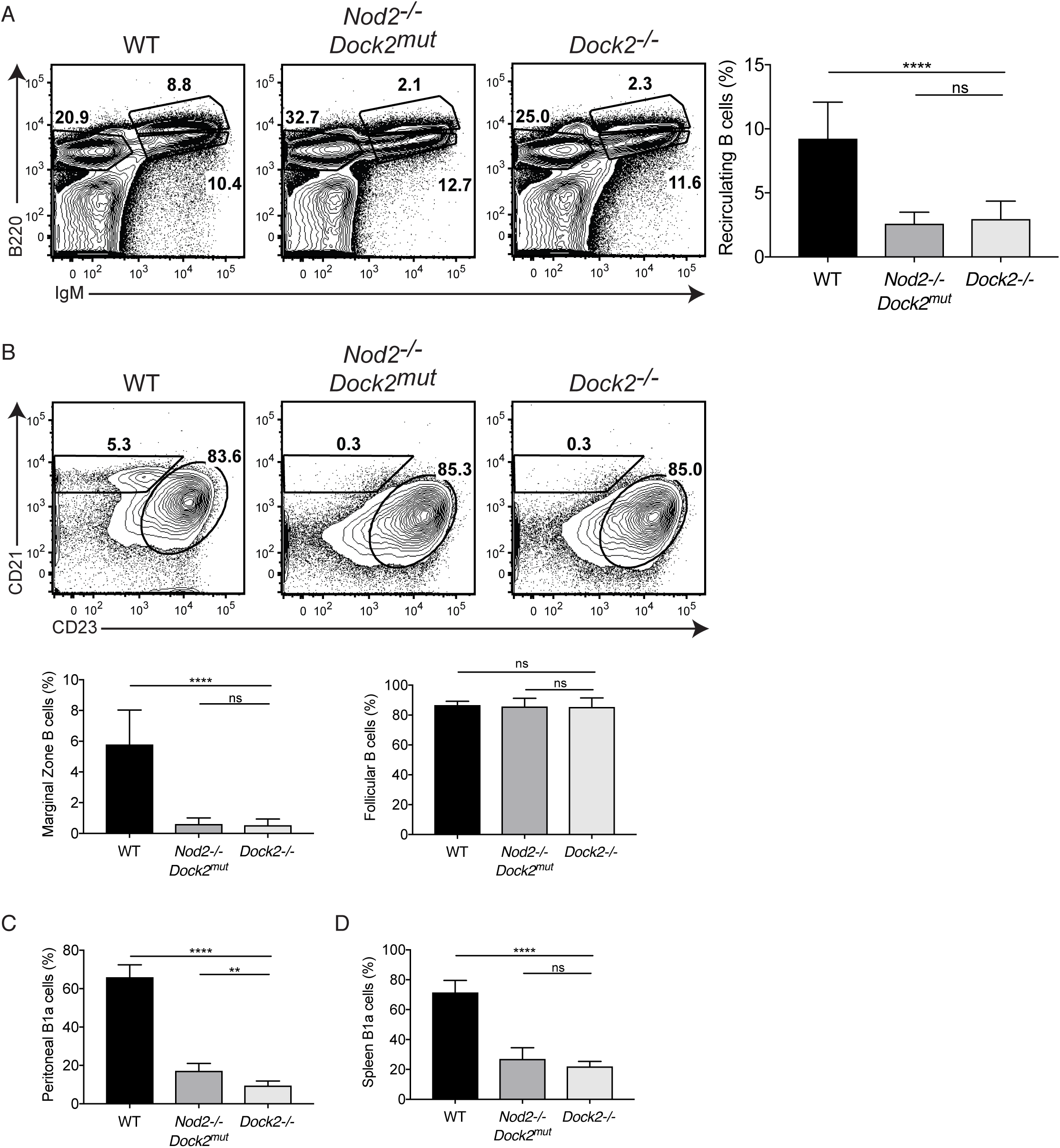
*Dock2*^−/−^ mice have defects in recirculating, marginal zone, and B1a B cells. Comparison of B cell populations in WT (*n* = 12), *Nod2^−/−^Dock2mut* (labelled as *Nod2^−/−^* in previous figures) (*n* = 6), and *Dock2*^−/−^ (*n* = 11) mice. **(A)** Representative FACS plot and percentage of bone marrow recirculating B cells (B220high IgM+). **(B)** Representative plot and percentages of marginal zone (CD21high CD23low) and follicular (CD21low CD23high) B cells in the spleen. Populations are gated on B220+ CD19+ AA4.1-mature B cells. **(C-D)** Representative plot and percentages of B1a cells (B220low, CD19+ CD5+) from peritoneal lavage (C) and in spleen (D). Data is representative of three independent experiments. ns *p* > 0. 05. \*\**p* ≤ 0.01, \*\*\*\**p* ≤ 0.0001 by two-tailed Mann-Whitney.

## DISCUSSION

Mutation of *NOD2* is one of the strongest risk factors for Crohn’s disease (37–39). NOD2 is well characterized as a bacterial sensor, but linking this molecular function with specific pathological outcomes has been challenging. We were eager to examine the B cell instrinsic role of NOD2 in light of the critical role of B cells, and antibodies they produce, in intestinal homeostasis. Furthermore, analyses of immunoglobulin (Ig) titers in IBD patients revealed a significant reduction in IgA (40), changes in IgA subclass distribution (41), and an increase in mucosal IgG directed against bacteria compared with healthy individuals (42). A central role for B cells in intestinal disease is further supported by the observation that individuals with common variable immunodeficiency, a primary immune deficiency that leads to decreased antibody production, develop an IBD-like disorder that includes villi blunting and intestinal inflammation (43–47). Together, these observations encouraged us to examine the B cell intrinsic role of this bacterial sensor.

The profound lymphocyte-intrinsic phenotype in *Nod2* deficient animals was rewarding, but the sheer magnitude of the difference in multiple B cell subsets, in the absence of any immunogen, was, nonetheless, surprising given that this strain of animals has been used by many laboratories around the world in the context of IBD studies and beyond (14, 20, 23, 31, 48–60). Unexpected phenotypes in analysis of gene targeted animal models can be a consequence of differences in the microbiota, genetic drift, or experimental assiduousness. We previously attributed goblet cell abnormalities in *Nod2^−/−^* mice to the presence of a specific member of the microbiota and were able to validate the results in mice deficient for RIP2, the downstream signal tranducer of NOD2 (24). The finding that the B cell deficiencies were not reproduced in *Rip2*^−/−^ prompted us to examine commercially-available *Nod2^−/−^* mice. The animals we obtained from Jackson Laboratories were derived from the same gene targeted founders as the animals within our colony, yet lacked the B cell defects we had observed. Numerous studies have established a direct role of microbial triggers in immune development (61). Microbial composition within strains and individual mice can vary greatly within the same facility (62). A prior analysis of *TLR* deficient mice (63) highlighted the importance of microbial history of the mice as the authors found that familial transmission of microbiota, rather than genetic loss of the pattern recognition receptors, directed the observed immune phenotypes. For this reason we undertook cross-fostering and extended cohousing experiments to control for changes in microbiota within our strains as a consequence of animal husbandry history. Ultimately, analysis of heterozygote matings to generate *Nod2^−/−^* revealed that a genetic factor was at play.

Finally, our findings with *Dock2^−/−^* mice confirmed that mutation in *Dock2* is sufficient to explain the B cell deficiencies we initially identified in *Nod2^−/−^* mice and also demonstrated that these defects were due to a loss of DOCK2 function. In addition to maintaining MZ B cells in the spleen, we have revealed additional roles for DOCK2 in the B cell compartment. All together, our findings indicate that DOCK2 is required for the maintenance of proper numbers of recirculating bone marrow cells, MZ B cells and B1a cells in the spleen, and peritoneal B1a cells. Although the near complete absence of these cells precludes biochemical analyses of molecular mechanism, previous studies have indicated that DOCK2 functions as a Rho guanine nucleotide exchange factor (GEF) involved in immune signaling and cell migration (3, 7, 64), which could explain the importance of this molecule for maintenance of dynamic B cell populations (65). Given the association between *DOCK2* mutations and immune-deficiency in humans, an important future direction will be to carefully examine the B cell compartment in patients for defects similar to those we have identified. Coincidentally, DOCK2 was recently identified as a key driver of gene expression patterns associated with IBD, and *Dock2^−/−^* mice are susceptible to intestinal injury (66). Therefore, a B cell specific function of DOCK2 in IBD pathogenesis may require consideration.

## ACKOWLEDGMENTS

We would like to thank Dr.Yoshinori Fukui of Kyushu University in Japan for the Dock2 -/-mice.

## FOOTNOTES

This work was supported by US National Institute of Health (NIH) grants R01 HL123340 (K.C., Mv.d.B.), R01 DK093668 (K.C.), R01 DK103788 (K.C.), R01 AI121244 (K.C.), R01 HL125816 (SBK), R21 AI124129 (SBK and S.S.), R21 AI110830 (SBK), Faculty Scholar grant from the Howard Hughes Medical Institute, Stony Wold-Herbert Fund, and philanthropy from Bernard Levine (K.C.); K.C. is a Burroughs Wellcome Fund Investigator in the Pathogenesis of Infectious Diseases; Feinberg Lymphoma Grant (SBK); Irma T. Hirschl Career Scientist Award (SBK), Colton Center for Autoimmunity Award (SBK) and Beckman Foundation Award (SBK).

